# Assessment of genome packaging in AAVs using Orbitrap-based charge detection mass spectrometry

**DOI:** 10.1101/2021.09.02.458670

**Authors:** Tobias P. Wörner, Joost Snijder, Olga Friese, Thomas Powers, Albert J. R. Heck

## Abstract

Adeno-associated viruses (AAV) represent important gene therapy vectors with several approved clinical applications and numerous more in clinical trials. Genome packaging is an essential step in the bioprocessing of AAVs and needs to be tightly monitored to ensure the proper delivery of transgenes and the production of effective drugs. Current methods to monitor genome packaging have limited sensitivity, a high demand on labour, and struggle to distinguish between packaging of the intended genome or unwanted side-products. Here we show that Orbitrap based charge detection mass spectrometry allows the ultra-sensitive quantification of all these different AAV bioprocessing products. A protocol is presented that allows the quantification of genome packed AAV preparations in under half an hour, requiring only micro-liter quantities of typical AAV preparations with ~10^13^ viral genome copies per millilitre. The method quickly assesses the integrity and amount of genome packed AAV particles to support AAV bioprocessing and characterization of this rapidly emerging class of advanced drug therapies.

## Introduction

AAVs are small viruses with a ssDNA genome that is encapsidated by a stochastic mixture of 60 VP1, VP2 or VP3 (*1*, *2*). The virus belongs to the genus *Dependoparvovirus,* and consequently is dependent on coinfection with helper viruses like Adeno- or Herpesviruses. AAVs inability to replicates on its own, combined with a low immunogenic profile make them ideal gene therapy vectors, with already several AAV-based gene therapy treatments approved by the EMA and FDA (*3*).

For gene delivery purposes, the natural AAV genome is replaced with a transgene, flanked by the natural AAV inverted terminal repeats (ITRs). These recombinant AAVs (rAAV) are typically produced in dedicated host expression systems where the transgene, the structural AAV genes and the required helper genes are co-transfected or stably integrated (*4*–*6*). Currently there are primarily two expression systems used, either human HEK293 or Sf9 insect cells. In either case, the produced rAAV particles must be purified from the cultured cell stock. Besides removing crude biological materials, this step also requires the proper separation of the empty from genome-filled AAV particles. The empty particles are still immunogenic but unproductive for transgene delivery, while accounting for up to 80% of the total number of particles produced in the cell culture system. Hence, just a fraction of total yield of AAV particles from the production system contains the desired genome (see Figure 1). This diversity of products, combined with the poor scalability of the used expression systems, makes AAV production and purification challenging and expensive.

**Figure 1:**
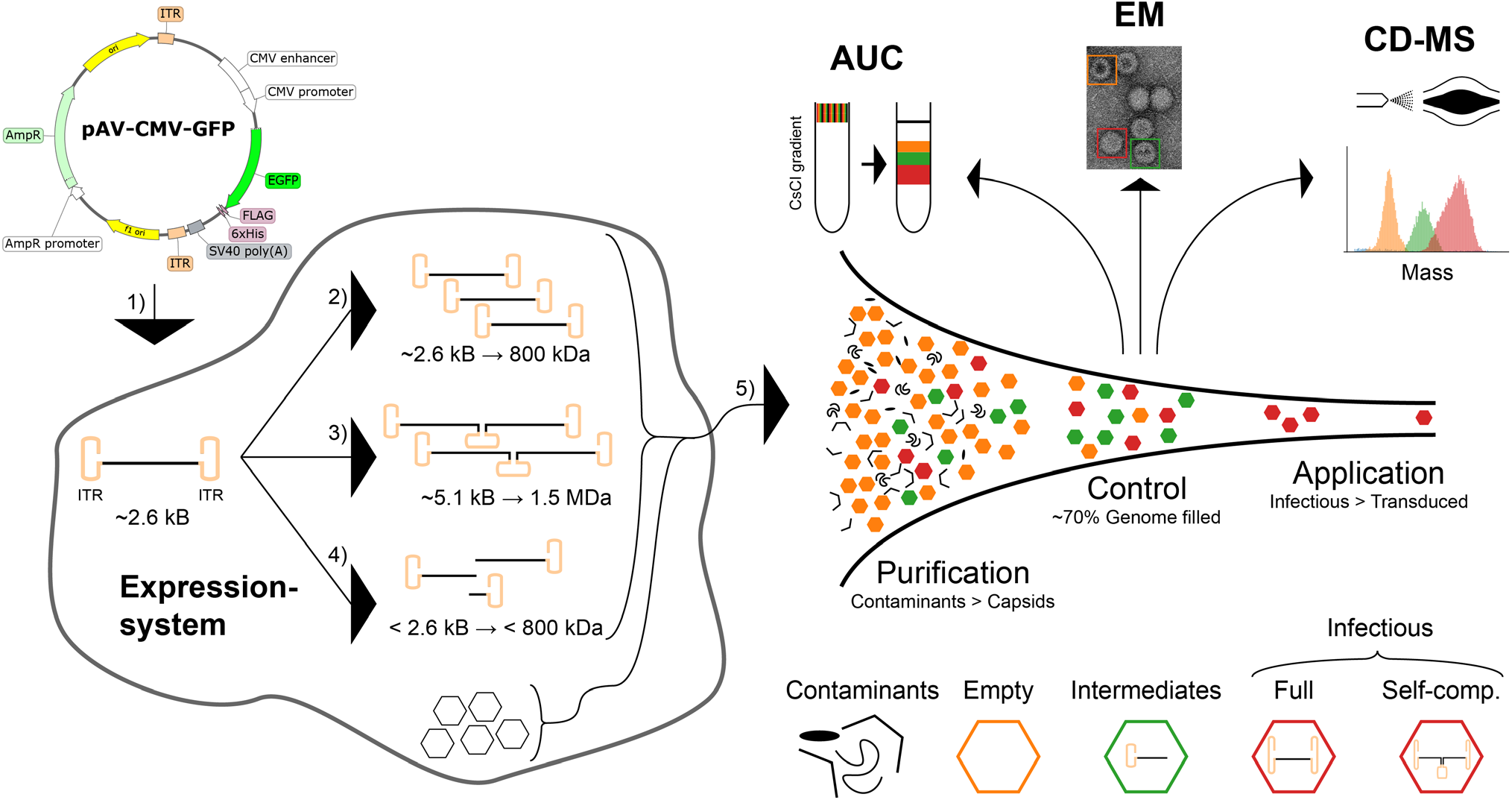
AAV bioprocessing and analysis. Schematic overview of rAAV production in either HEK293 or SF9 host cells for a designed transgene of around 2.6 kb. After transfection of (or infection with) the corresponding transgene (1) (if replicated properly) will yield an encapsulated 800 kDa genome (2). Other encapsulated off-target genomes could originate from self-complementary ssDNA dimeric variants (3) as well as truncated genomes (4). These gene products are all encapsulated (5) and will yield a mixture of empty and (partly) filled particles. The initial number of other contaminants (DNA, side product, host cell proteins) can be several orders of magnitude higher than that of rAAV capsids and require extensive purification. rAAV capsids show additionally a wide distribution of empty, partially filled, and filled capsids, requiring additional purification and monitoring. Detailed characterization is important as only a small number of the infectious particles will transduce their genome. This efficiency is sensitive to the presence of empty capsids and capsids filled with off-target genomes. For monitoring this, AUC and EM are the industry standards, whereby the former can also serve as preparative method. Here Orbitrap based CD-MS is explored for quality control, as it is sensitive, facile and can yield information on capsid/genome integrity.

Ideally, the filled AAV particles will only contain the ITRs flanked genome. However, several processes may lead to genome heterogeneity resulting in capsids partially filled with truncated genomes or even particles that package multiple genome copies (see Figure 1). First, if the packaged transgene exceeds the natural genome capacity of ~4.8 kb the genome may become truncated during the encapsidation (*7*). Such AAV preparations, similar as in dual-vector strategies, still can transduce the corresponding gene of interest (GOI) via homologous recombination in the target cell, albeit at a greatly reduced efficiency (*8*, *9*). Second, genome heterogeneity can also already be induced during replication. Like all *Dependoparvoviruses*, AAV genome replication follows the “rolling hairpin” model where the Rep endonuclease nicks the terminal resolution site (*trs*) to allow the synthesis of the second downstream ITR (*10*). If this process fails, the final product can be a self-complementary single strand genome at approximately twice the size of the monomeric DNA (*11*). This process can be exploited by modifying the Reb binding element (RBE) to promote the formation of self-complementary AAV (scAAV), but this process occurs also at a low rate in wt-ITRs (*12*–*14*). Administration of scAAV have the benefit that transcription can start directly from the internally hybridized dsDNA transgene and does not require the rate limiting synthesis of the complementary DNA strand and intermolecular hybridization (*15*, *16*). Third, secondary structure elements in the DNA can cause genome heterogeneity as has been documented in studies whereby rAAV had been designed to deliver short hairpin RNAs and CRISPR elements, which were shown to interfere with the genome replication (*17*, *18*). This latter feature is especially important in targeted gene delivery, as many mammalian transgenes contain highly structured elements to assist in promotor binding.

Sensitive and specific methods to investigate the processes leading to unwanted genome heterogeneity may lead the way to improved bioprocessing of homogeneous AAV particles for clinical applications. Typical lab scale preparations of rAAVs yield no more than a few milliliters of purified samples (~10^12^-10^14^ viral genomes in total), which makes subsequent analysis challenging, especially when screening several different experimental growth conditions. From the biophysical methods available to investigate potential heterogeneity in rAAV preparations, analytical ultracentrifugation (AUC), electron microscopy (EM) and polymerase chain reaction (PCR)-based methods are currently the industry standards (see Figure 1), but all these approaches have also their downsides (as reviewed in (*19*)). While AUC is also used as a preparative purification method to fractionate empty from filled rAAV particles, it requires relatively large amounts of samples (0.5 ml of ~2*10^12^-5*10^12^ vc/ml) and while the resolving power is adequate to separate empty from filled particles, it is challenging to dissect particles filled with distinct genome intermediates. Negative stain EM requires typically much less sample but staining and drying of the samples may disturb particle integrity leading to genome release and a bias towards empty capsids. Moreover, these staining artefacts further complicate distinguishing fully packed capsids from the partially filled side-products by EM. PCR-based methods require just minute amounts of sample, but mostly can only report on genome titer, without distinction of genome integrity. Thus, currently, the field is missing a quantitative method to distinguish between full, empty, and partially packaged rAAVs with sufficient resolution, sensitivity and dynamic range to detect even the lowest abundant populations. Moreover, methods that can also be used to characterize the nature of these potential genome intermediates, for instance by measuring their mass or sequence their DNA, would be useful for optimizing the rAAV production processes.

A relatively new alternative approach used for rAAV characterization is charge detection mass spectrometry (CD-MS) as first demonstrated by using home-build instruments (*20*). The feasibility to perform CD-MS on commercial Orbitrap UHMR platforms was recently demonstrated and also used for analyzing rAAVs (*21*, *22*). In CD-MS particles are measured individually as opposed to conventional native MS. Inherently, single particle measurements offer extreme sensitivity and thus low sample consumption, which is highly beneficial for successful application to clinical preparations of rAAV.

Here we present an optimized workflow enabling the mass analysis of heterogeneous rAAV particles, whereby the mass resolving power and accuracy attainable allow us to determine and quantify genome integrity. We determine the mass and abundance of various co-occurring rAAV particles in less than 30 minutes, using minimal amount of sample. We introduce an improved acquisition method with better signal utilization, that can accurately and reproducible detect rAAV particle populations as low as 2%, also identifying some lower abundant, likely scAAV variants, as off-target products in rAAV preparations. We show that the quantitative accuracy of the CD-MS method to distinguish filled from empty particles is 1-2%. This new approach can contribute to the optimization of the bioprocessing of rAAVs, producing more particles containing the desired intact transgene.

## Results and Discussion

### Optimizing the analysis of co-occurring AAV particles in CD-MS

Here we aimed quantitatively measure co-occurring rAAV particles ranging in mass from about 3.6 (empty capsids) to 5.3 MDa (genome packed capsids). Toward this goal we build further on our previous study of AAV by Orbitrap-based charge detection mass spectrometry (*22*). In this approach rAAV particles are directly diluted, from their storage solution, in aqueous ammonium acetate and introduced into a mass spectrometer where they are ionized by electrospray ionization under native conditions (*23*). Subsequently each particle is individually detected within the Orbitrap mass analyzer and we can estimate the number of charges for each ion directly from its intensity. This enables mass determination on a single-particle basis with the particle counts providing quantitative information on population distributions. The unbiased detection of particles of different composition and mass is not trivial in MS as ion transmission and ion decay processes are charge and mass dependent (*24*, *25*). Therefore, we initially investigated the effect of the pressure settings and transient recording times on the detection of all different co-occurring rAAV particles and observed that transient times of 512 ms, and pressure settings between 1.5 and 3, provided the optimal conditions for the least biased detection of all co-occurring particles, as described in detail in the Supplementary Materials and Supplementary Figure 1.

### Assessing AAV particle diversity

The importance of using this optimized acquisition approach becomes most eminent when aiming to analyze and quantify low abundant genome variants as demonstrated in Figure 2. The analyzed particles were expressed with a common expression cassette with an approximately 2.5 kb long genome (with a Mw of ~800 kDa). Besides packing this genome, other possible genome variants that may be packed by the AAV are a self-complementary ssDNA (Mw ~2*800 kDa) and truncated forms of the genome as further depicted in Figure 1. By CD-MS, the obtained 2D histogram and the corresponding mass histogram for an AAV6a serotype, expressed with the above-mentioned transgene are shown Figure 2A and 2B. Cumulatively, the 2D CD-MS spectrum shown in Figure 2A shows the data for about ~76,000 individual rAAV6a particles. From 2D CD-MS two prominent co-occurring distributions can be observed, but also two weaker particle populations, all with distinct masses. The two most prominent particle populations represent the empty (~23,000 particles, Mw = 3.85 MDa) and genome filled rAAV6a particles (~40,400 particles, Mw = 4.63 MDa), with a relative mass difference of ~800 kDa, due to the packaging of the expected genome. All measured masses are summarized in Table 1. The additional densities in the 2D mass histogram appearing between the “empty” and “full” as well as at higher *m/z* beyond the “full” particles, exhibit masses of approximately 4.3 MDa (~3,700 particles) and 5.3 MDa (~1,800 particles), respectively. The particle population with the highest mass likely represents AAV capsids packaging a self-complementary transgene version. The intermediate mass ions are likely AAV capsids encapsulating truncated transgenes and show a less clearly defined mass distribution (see also Supplemental Figure 2). The mass difference (relative to the empty capsid) of these lower mass particles correlate quite well with the location of G/C rich structures in the EGFP sequence, already identified as a putative source for likely genome truncation (*17*). Notably, all these co-occurring particles can be resolved in the 2D CD-MS spectra, whereby also the dynamic range reached between the most abundant and least abundant particle distribution is close to 40, with the rAAV6a particles seemingly packaging dimeric DNA representing just 2.4% of all particles, and the rAAV6a particles packaging truncated DNA representing just 4.8% of all particles.

**Figure 2:**
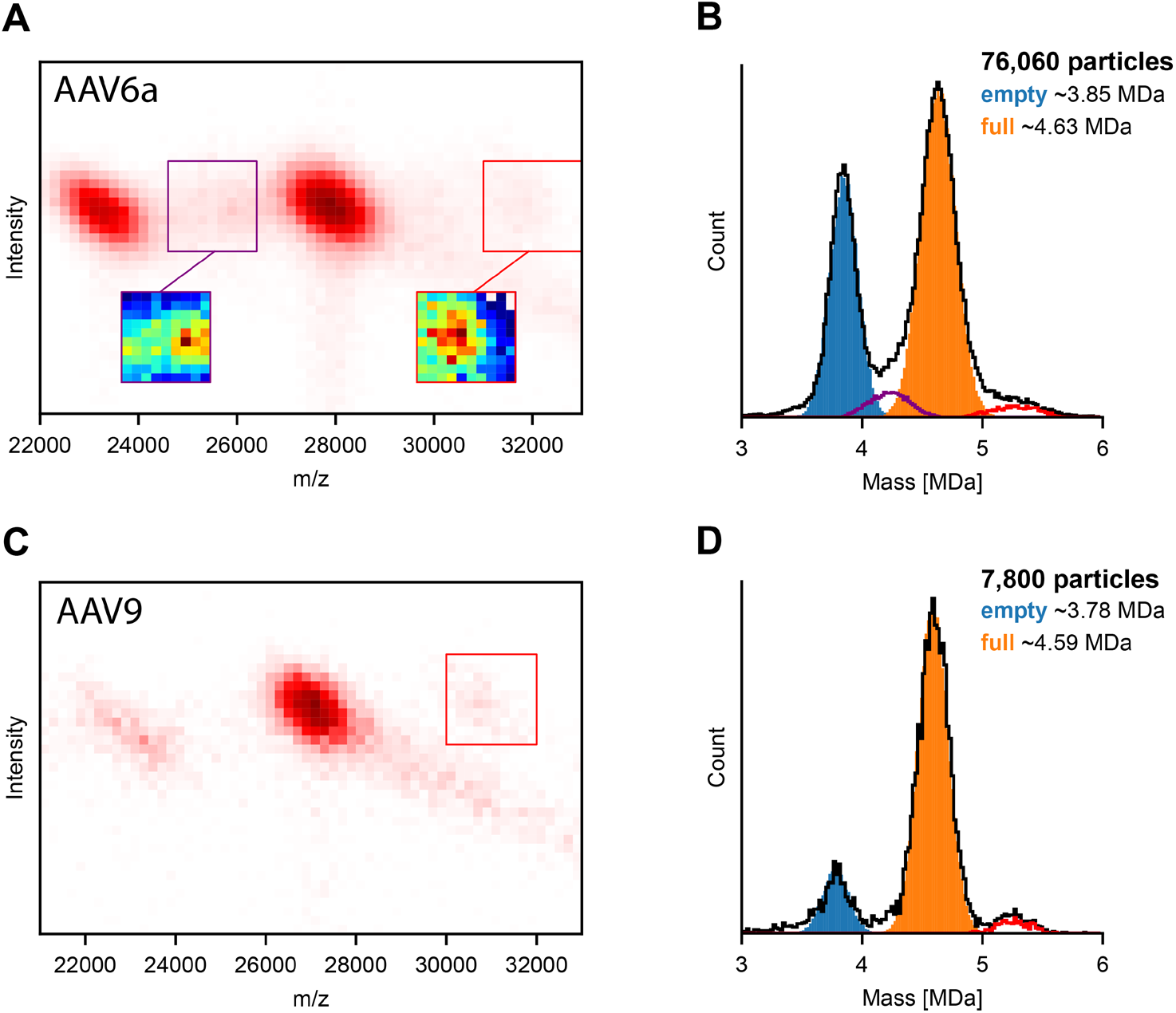
AAV bioprocessing can yield a diversity of co-occurring particles that can be sorted and counted by CD-MS. **A)** 2D CD-MS histogram of an AAV6a preparation revealing the mass and abundance of co-occurring particles and in **B)** the corresponding mass histogram. Besides the distributions originating from the empty and fully filled particles, low-abundant signals are observed in the 2D histograms corresponding to particles containing the self-complementary genome variants (red box) and intermediate truncated variants (purple box). The particles contained within these latter boxes are shown in the in the mass histogram with an alike color scheme. **C)** 2D CD-MS histogram of an AAV9 preparation and in **D)** the corresponding mass histogram.

**Table 1:** Measured and estimated expected masses for distinctive type of particles co-occurring in the preparations of rAAV6a, rAAV9 and rAAV6b. The measured experimental masses correspond to the average masses determined from the mass histograms extracted from the CD-MS data. The expected masses are based on the amino acid sequences of the AAV VP proteins and an assumed stoichiometry of 60 capsid proteins (assumed 5:5:50 for VP1:VP2:VP3 and with ±5 VP as composition can vary (*1*)), while the mass of the DNA is estimated from its nucleic acid sequence. For the AAVs of serotype 6 the AAV6a particles were expressed with a GFP genome, whereas the AAV6b were expressed with a proprietary genome.

| rAAV6a | Measured [MDa] | Expected [MDa] | Genome [kbp] | Genome [MDa] | Assigned |
| --- | --- | --- | --- | --- | --- |
| Empty | 3.85 | 3.71±0.1 | - | | Empty particles |
| Filled 1 | 4.63 | 4.65 | 2.6 | 0.8 | Filled particles |
| Filled 2 | 5.30 | - | 5.1 | 1.6 | Dimeric genome |
| Filled 3 | 4.0-4.4 | - | 1.0-1.5 | 0.15-0.55 | Truncated genome |
| rAAV9 |  |  |  |  |  |
| Empty | 3.78 | 3.72±0.1 | - | | Empty particles |
| Filled 1 | 4.59 | 4.58 | 2.6 | 0.8 | Filled particles |
| Filled 2 | 5.2 | - | 5.1 | 1.6 | Dimeric genome |
| rAAV6b |  |  |  |  |  |
| Empty | 3.7 | 3.71±0.1 | - | | Empty particles |
| Filled 1 | 5.2 | 5.3 | 5.1 | 1.6 | Filled particles |
| Filled 2 | 4.4 | - | 2.5 | 0.8 | Truncated genome |

To further validate that these lower abundant populations of AAV particles are observed also in other rAAV preparations and other rAAV serotypes, we next analyzed by CD-MS a rAAV9 sample, expressed with the same transgene. The CD-MS-obtained 2D histogram and the corresponding mass histogram for an AAV9 serotype are shown Figure 2C and 2D. Also in this 2D CD-MS two co-occurring distributions are observed, originating from the empty (~1,000 particles, Mw = 3.78 MDa) and genome filled rAAV6a particles (~5,600 particles, Mw = 4.59 MDa), but again a lower abundance particle population is observed, for rAAV9 particles containing likely dimeric DNA.

### Assessing AAV particle abundances

With the above-described improved data acquisition method, we developed a dedicated workflow for the sensitive and accurate mass analysis of rAAV preparations, and next wanted to assess its performance especially in the quantification of co-occurring particle distributions. The aim here was to minimize sample consumption as well as demands on time and labor (see Figure 3) to meet industry standards. Based on its improved sensitivity and direct charge assessment CD-MS is less negatively affected by high salt concentrations present in the storage buffer and consequently we observed that the rAAV samples could be directly diluted into aqueous ammonium acetate instead of requiring a time- and sample consuming buffer exchange by dialysis or spin filters. We found for our AAV preparations (2*10^13^ vc/ml) a dilution factor of 20 sufficiently reduced the salt concentration, leaving a high enough AAV concentration for fast data acquisition. We were able to measure and record around 30,000 particles in 20 minutes, which is sufficient for good statistics in the down-stream data analysis. The whole sample preparation as well as data acquisition could be performed in less than 30 minutes. The data processing pipeline is largely automated and can be, after initial setup, executed with a runtime of less than a minute per dataset.

**Figure 3:**
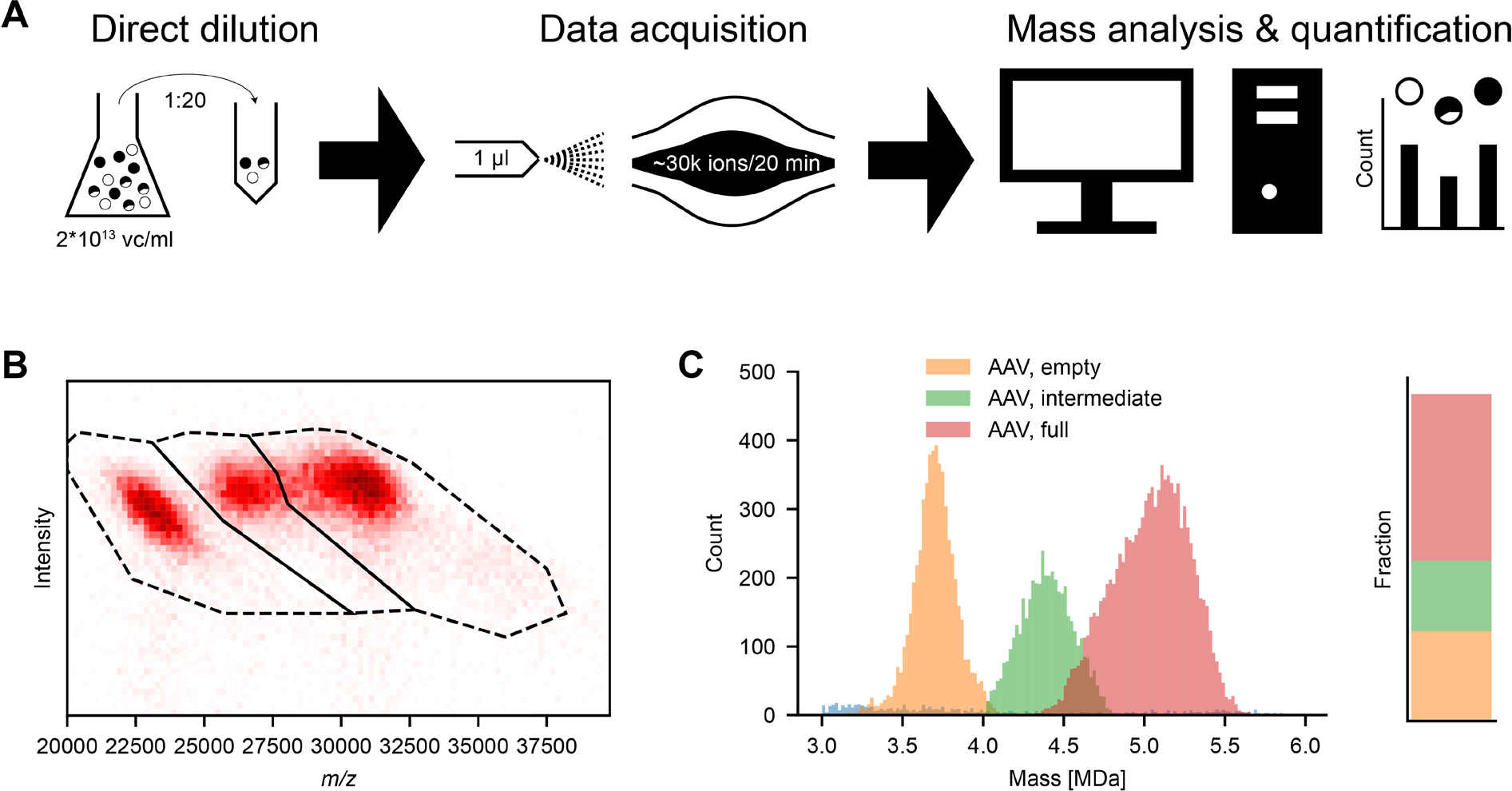
Direct sampling enables efficient AAV particle characterization and quantification by CD-MS. **A)** Optimized sensitive approach for AAV particle characterization and quantification. Speed and required input amount is significantly reduced as no buffer exchange is needed. Here, the depicted workflow can be executed in ~30 minutes per sample. **B)** Illustrative 2D histogram of a dataset acquired in 20 min. For fast and easy quantification, all ions which fall in the margins of the drawn polygon are either counted as empty, partially full or full AAV particles. **C)** Corresponding mass histogram and bar plot depicting the most abundant masses for each species as well as their relative abundance.

Establishing this sensitive and efficient workflow we focused for our quantitative analysis on an industrial AAV6 sample with a proprietary transgene, that we term here AAV6b, to distinguish from the commercial AAV6a sample. The produced AAV6b particles were first separated into full and empty particles by using a CsCl gradient centrifugation. Following this fractionation, the empty and full rAAV particles were mixed in defined ratios of 0, 25, 50, 75 and 100%. These 5 distinct samples were subsequently analyzed by CD-MS. The setup as well as an illustrative example of the recorded data are visualized in Figure 3B and 3C for a rAAV6b sample, in which the empty and full fractions had been pre-mixed in a ratio of 25%:75%. However, for this rAAV6b sample we observed in the 2D CD-MS histogram clearly three distinct particle distributions which we assigned, based on their mass, to empty, intermediate and full genome packed rAAV capsids. We constructed for each observed distribution in the 2D CD-MS spectrum a polygon capturing the majority of the corresponding ions, used to filter and assign all ions laying within the polygons margins. From the filtered ion distributions, we can calculate for each particle population the average mass as well as count the particle population, as presented in Figure 3C. For this particular rAAV6b sample we assigned a capsid mass of 3.7 MDa (5,651 particles, orange), a mass of 5.2 MDa (10,529 particles, red) for the full genome packed AAV particles and 4.4 MDa (4,490 particles, green) for AAV particles packaging a truncated genome, respectively. The quantification based on these particle counts yields a ratio of 27:22:51% (empty:partially full:full) for this sample, which is in good agreement with the fact that this sample had been pre-mixed with 25% empty particles. However, our data reveal that a substantial part of the seemingly filled rAAV6b particles, fractionated and purified by using analytical ultracentrifugation, are not filled with the desired transgene, but just a truncated part of that.

To further evaluate the method, we next measured samples from the wider range of mixing ratios, also under two distinct experimental conditions, varying the pressure settings (see Figure 4A and Supplemental Figure 3) to further confirm that they do, in the applied pressure range, not influence the qualitative and quantitative outcome of the analysis. The resulting 2D CD-MS data and mass histograms are shown in Figure 4A, with from left to right the empty to completely “full” rAAV particles. For the quantitative analysis the same three polygons, and color-coding, were used to count the particle distributions. The relative contributions of each of the three distinct particle populations in the 5 measured pre-mixed samples are depicted in Figure 4B, whereby the two bars shown for each sample represent the measurements at the two different experimental pressure settings. Firstly, this analysis reveals that in the used pressure regime, the change in pressure does not affect the particle distribution. Secondly, when summing up all the particles packing the intact and truncated transgenes, and considering them as “full”, we observe that the ratio between empty and full rAAV as measured by CD-MS, matches very well the expected ratio, based on the known mixing ratio. The expected ratio is depicted by the dashed black line in Figure 4B. Although the by CD-MS assessed ratio is already pleasingly in agreement with the expected ratio, we noticed that in the measurements of the pre-mixed samples there were slightly more empty particles detected by CD-MS than expected based on the mixing ratio. To investigate this slight discrepancy, we carefully examined also the CD-MS data of the non-mixed empty and full samples. While the empty sample seems to be relatively pure, we still observed empty particles in the presumable pure full fraction (Figure 4A and 4B). This contamination of empty rAAV particles is likely caused by the insufficient resolving power in the ultracentrifugation. Measuring the pure “full” sample we could assess the percentage of partially and fully filled particles, and observed that in this particular sample there is a relatively high amount of particles that seemingly have packed a truncated DNA molecule (about 26% of all filled particles) as shown in Figure 4C. A similar 30%:70% ratio between half full and full rAAV was consistently measured also after mixing the full fraction with an empty fraction, as shown in Figure 4C. Evidently, this is to be expected but also shows that the quantification of particle populations by CD-MS can be quite robust and accurate, using only minute amounts of sample and time.

**Figure 4:**
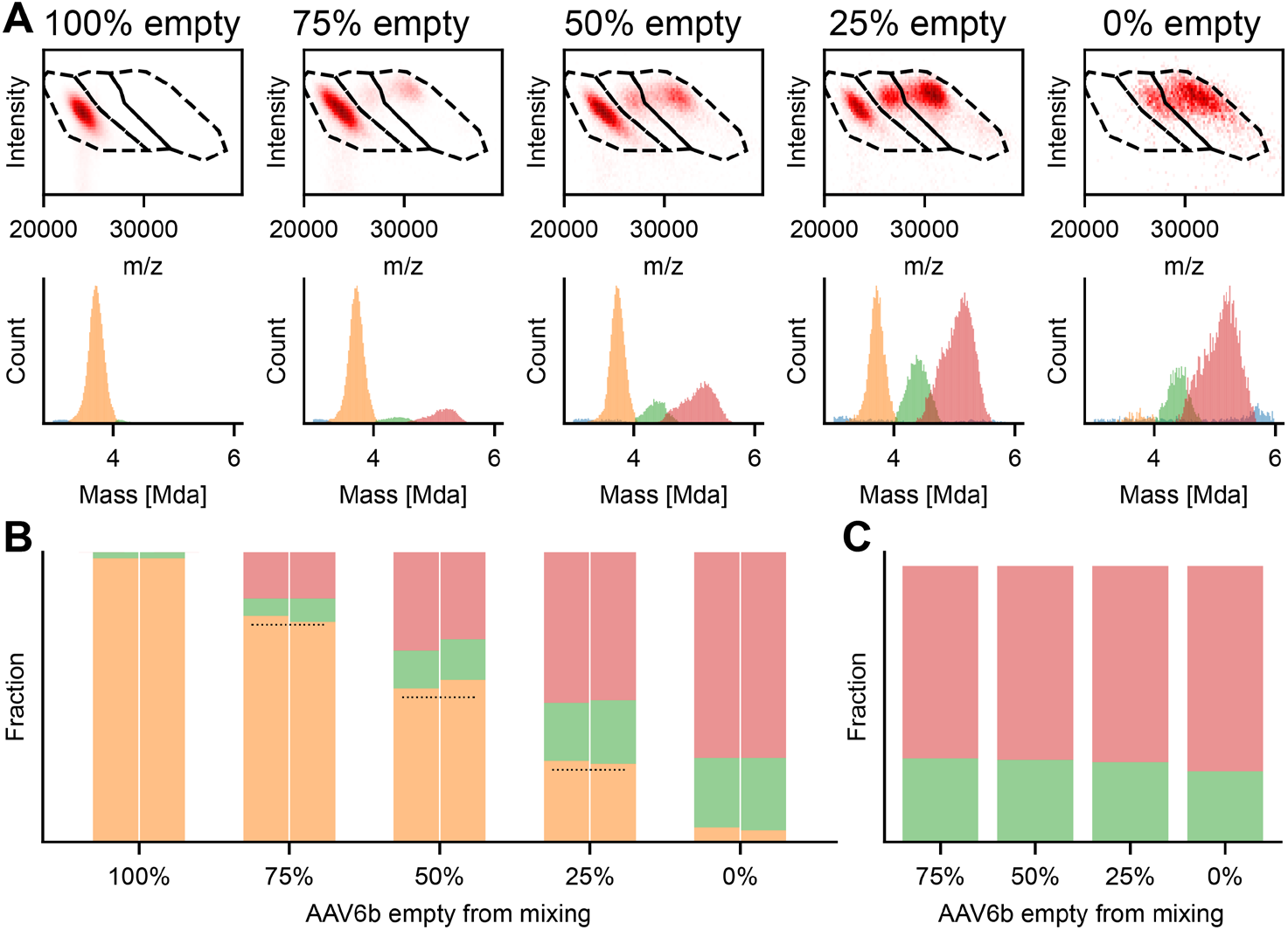
Quantification of distinct rAAV6b particles. **A)** 2D histograms (top row) and corresponding mass histograms of AAV particles mixed at defined ratios (the estimated % of empty particles is indicated above). **B)** Fractional abundances of empty, partially and fully filled AAV particles from **A)** and Supplemental Figure 3a. Dotted lines indicate the expected estimated values of 75%, 50% and 25%. For each mixing ratio, CD-MS data gathered at two gas pressure settings are shown (Xe 2 and 3, respectively on the left and right). **C)** The relative fractional abundance of the partially and fully filled AAV particles, remains constant with about 71% fully packed and 29% packed with a truncated genome.

## Materials and Methods

### AAV samples

Full and empty mixture of AAV6a and AAV9 were purchased from Vigene Biosciences containing a GFP genome. The alternate AAV6b samples were produced by Pfizer using a baculovirus expression system containing a proprietary genome.

For the AAV6b samples the enriched empty and full particles were fractionated via cesium chloride ultracentrifugation using a Beckman Coulter Optima L80XP ultracentrifuge with a SW 41Ti swinging bucket rotor run at 40,000 rpm for 44 hours at 15°C. Following centrifugation, two distinct viral bands were visible: a low-density and high-density band, corresponding to empty and full capsids, respectively. Each purified band was collected via side puncture and buffer exchanged into PBS using a 10kDa molecular weight cutoff dialysis devise. The capsid concentration of each bulk enriched fraction was determined by size exclusion chromatography, followed by subsequent dilution in a proprietary formulation buffer at equivalent sample concentrations. The theoretical percentage of empty capsids assumes that the low-density band is 100% empty capsids and that the high-density band is 100% full capsids.

### AAV CD-MS

For CD-MS analysis 1 ul of 2E13 vp/ml were diluted directly in 19 ul aqueous ammonium acetate (75 mM, pH 7.5) and subjected to single particle CD-MS analysis. Therefore, an aliquot of 1 μl was loaded into gold-coated borosilicate capillaries 467 (prepared in-house) for nano-ESI. Samples were analyzed on a standard Q Exactive UHMR instrument (Thermo Fisher Scientific, Bremen, Germany) (*25*, *26*). The instrument parameters were optimized for the transmission of AAV particles. In short, ion transfer target *m/z* and detector optimization were set to ‘high *m/z*’. In-source trapping was enabled with desolvation voltage of −150. The ion transfer optics (injection flatapole, inter-flatapole lens, bent flatapole and transfer multipole) were set to 10, 10, 4 and 4. We used Xenon as neutral gas in the collision cell and complexes were desolvated via activation in the HCD cell. Gas pressure was varied during experiments and the mentioned gas pressures correspond to UHV cold cathode gauge after applying the correction factor. Data was recorded at either 512 or 1024 ms transient time as specified in the manuscript.

### AAV quantification

For AAV quantification acquired .raw files were converted to mzXML with vendor peak picking enabled using msConvert. If not mentioned otherwise, data files were used for quantification without filtering. If filtering was applied, first, all centroid above the noise level were removed, then split peak occurrences were removed by removing each peak with a neighboring peak within 3 times the FWHM of the theoretical achievable resolution. Ions were quantified either based on their *m/z* position or by fitting a Gaussian distributions into the mass histograms.

## Author Contributions

T.P.W., J.S., and A.J.R.H. conceived the project, designed the experiments, and wrote the paper. O.F. and T.P. provided the fractioned AAV6b samples. TPW performed and analyzed the single particle CD-MS experiments. J.S. and A.J.R.H. supervised the project. All authors discussed the results and edited the paper.

## Conflicts of Interest

O.F. and T.P. are employees of Pfizer WRDM, St Louis, MO, a company with interest in employing AAV vectors for gene delivery purposes. The remaining authors declare no competing interests.

## Acknowledgements

We thank the members of the Heck laboratory for general support, especially Arjan Barendregt. This research received funding through the Netherlands Organization for Scientific Research (NWO) through the Spinoza Award SPI.2017.028 to A.J.R.H. Additionally, we are grateful for the support from Pfizer Biotherapeutics Pharmaceutical Sciences. Specifically, Qin Zou and Cortney Sloan are acknowledged for their project support and generation of orthogonal data for baseline characterization of AAV6b material.

## Supplementary Text

### Optimizing conditions for the quantitative analysis of rAAV particles; pressure dependence

The pressures in different sections of a mass spectrometer are important parameters in native MS experiments and must be balanced delicately to ensure a successful experimental outcome (*1*–*3*). Although generally high vacuum conditions are important in mass spectrometry, a somewhat elevated pressure, allowing collisional cooling, enhances effective transmission of high mass ions species through the mass spectrometer (*4*, *5*). On the other hand, collisions with background gas can cause ion decay, and lead to eFT artefacts resulting is peak splitting in the acquired signals causing a systematic trailing to lower intensities in Orbitrap based CD-MS (*6*, *7*). We illustrate this effect in Supplemental Figure 1A, where we used increasing gas pressure settings and investigate its effect on the transmission of different AAV6a capsid variants. As more often used in native MS we introduced Xenon as collision- and cooling gas. While for the lowest pressure setting, we observed barely any trailing to lower intensities, the transmission of the filled rAAV capsid particles at 28,000 *m/z* became significantly reduced. However, at elevated pressure settings, the transmission of the full rAAV species increased but at the same time, the frequency of peak splitting increased as well, requiring an additional filtering step in the downstream data analysis. The results of this peak splitting and filtering on the final mass histograms are depicted in Supplemental Figure 1B. In the unfiltered data, we observe some density trailing to lower masses (at around 3 MDa) associated with a reduced peak intensity of the split peaks. These “low-mass”-artifacts increase the FWHM of the corresponding distribution and cause a slight underestimation of its corresponding average mass. Additionally, as higher mass ions seem to be more prone to peak splitting, such centroids will be removed more frequently in the signal filtering step, biasing the quantification of filled and empty capsids and furthermore, reducing sensitivity for high mass, low abundant ion species. These artifacts can be mitigated by reducing somewhat the transient recording time from 1024 to 512 ms, as demonstrated previously (*8*). The effect of a shortened transient time on the mass histogram is also shown in Supplemental Figure 1C, where we can see little to no difference between the unfiltered and filtered datasets, when recording for 512 ms. This allows us to access a much wider pressure range which we examined for the ratio of full and empty as well as potential top mass reduction as illustrated in Supplemental Figure 1D. From these analyses we define a Xenon pressure setting between 1.5 to 3 to give the best balance between optimal and even transmission of all rAAV particles, reducing as much as possible artefacts induced by ion decay processes due to interactions with the neutral gas molecules. Therefore, we decided to use a Xenon pressure setting of 2 and 3 for the analysis of the rAAV preparations. The conversion for the used pressure setting to the resulting cold cathode gauge readout are shown in Supplemental Figure 1E.

## Supplementary Figures

**Supplemental Figure 1:**
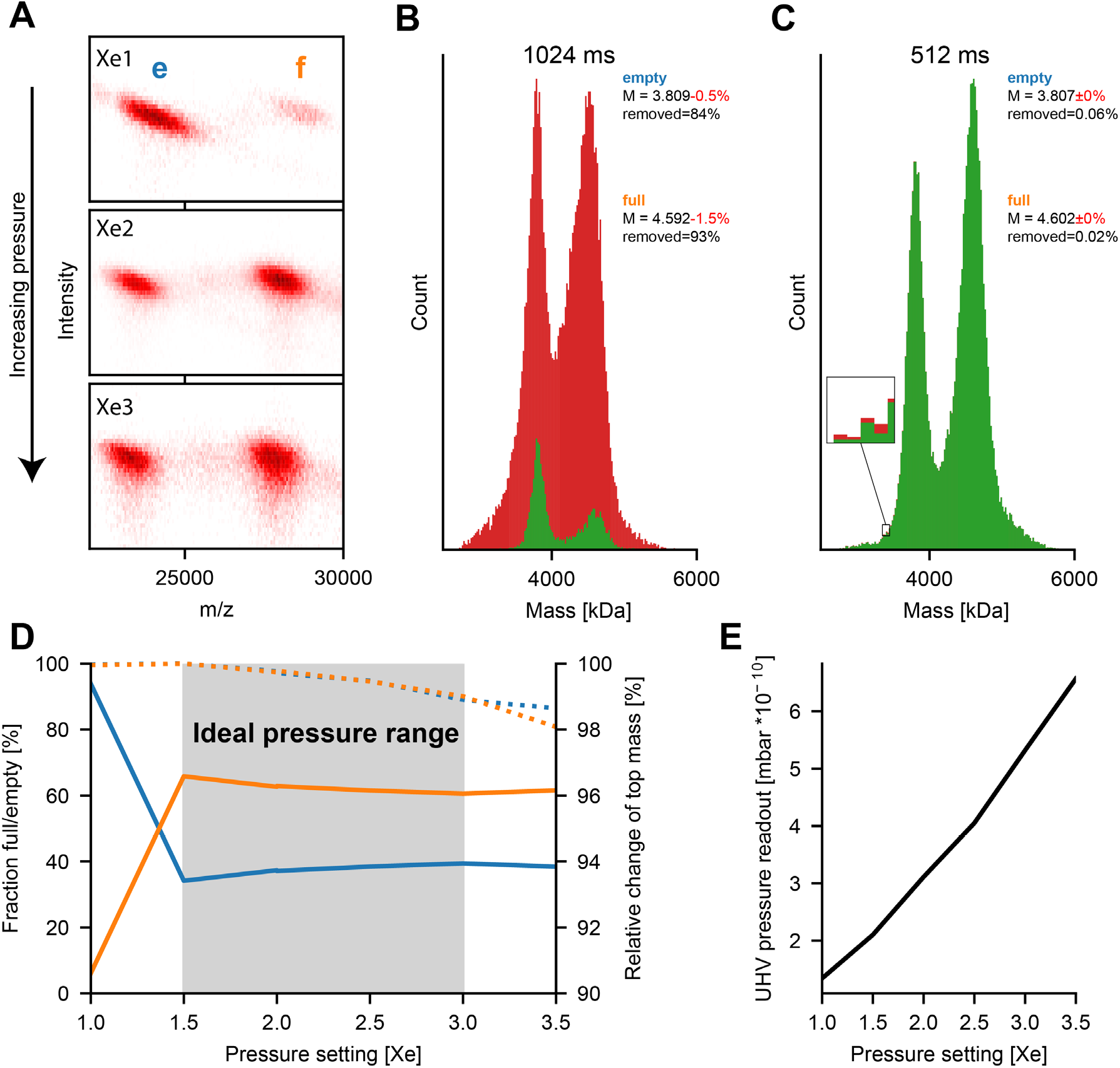
Optimizing pressure and transient times for the unbiased and sensitive analysis of co-occurring rAAV particles. **A)** 2D histogram of mixture of empty (**e**) and filled (**f**) rAAV capsids recorded at different pressure settings, illustrating the effect of background gas on the transmission as well as peak splitting. While the filled particles are hardly transmitted at lower pressure, compared to the empty species, elevated background pressure offers a flatter transmission profile but also increased peak splitting at transient times of ~1024 ms. **B and C)**, Influence of split peak filtering on AAV capsid mixtures recorded at different transient times of 1024 ms and 512 ms (unfiltered and filtered mass histogram are shown in red and green respectively). The number of removed centroids as well as the assigned mass, including the change with respect to the unfiltered dataset, are indicated for each distribution. For the longer transient time (**B**) extensive peak splitting takes place, resulting in significant trailing towards lower masses for the unfiltered data (red). This trailing causes a reduced effective resolving power as well as a slight underestimation of the assigned masses. The required filtering recovers the suppressed resolution and top mass but removes a significant portion of the recorded data and suppresses low abundant signal, especially in the higher mass range (>5MDa). For the sorter transient time peak splitting is reduced significantly removing the need for filtering in general. **D)** Tested pressure range for optimal transmission of full and empty particles at ~0.5s transients time without filtering. Pressure setting between 1.5 and 3 show flat transmission profile for empty and filled particles while still only exhibiting insignificant amounts of split peak artifacts (reduced top mass). **E)** Relation between experimental pressure setting and UHV read-out pressure.

**Supplemental Figure 2:**
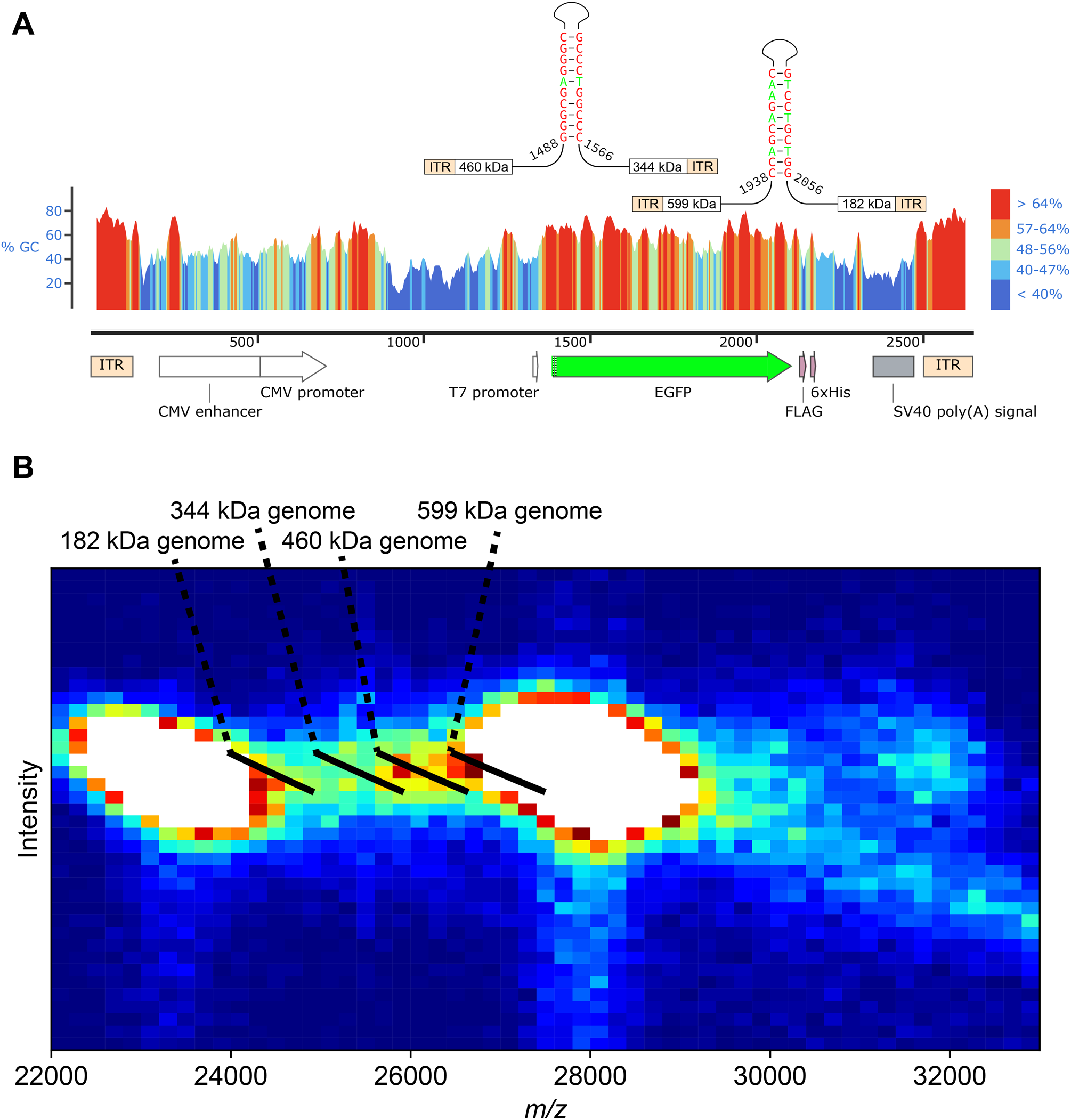
Detection of low abundant rAAV variants caused by heterogeneity in the packed DNA. **A)** The transgene cassette to be packed into a rAAV exhibits several G/C-rich (red) regions attributed to ITRs but also in the CMV promoter as well as in the EGFP sequence. The EGFP transgene contains particular regions of palindromic G/C-rich sequences where genome replication might be impaired. The expected genome weight is indicated for a replication termination of the corresponding sense or antisense strand. **B)** 2D Histogram taken from Figure 2A with altered color coding emphasizing the occurrence of low abundant species. The expected position of a capsid encapsulating the potential truncated genomes in **A)** coincide with the detected signals between the empty and filled rAAV particles.

**Supplemental Figure 3:**
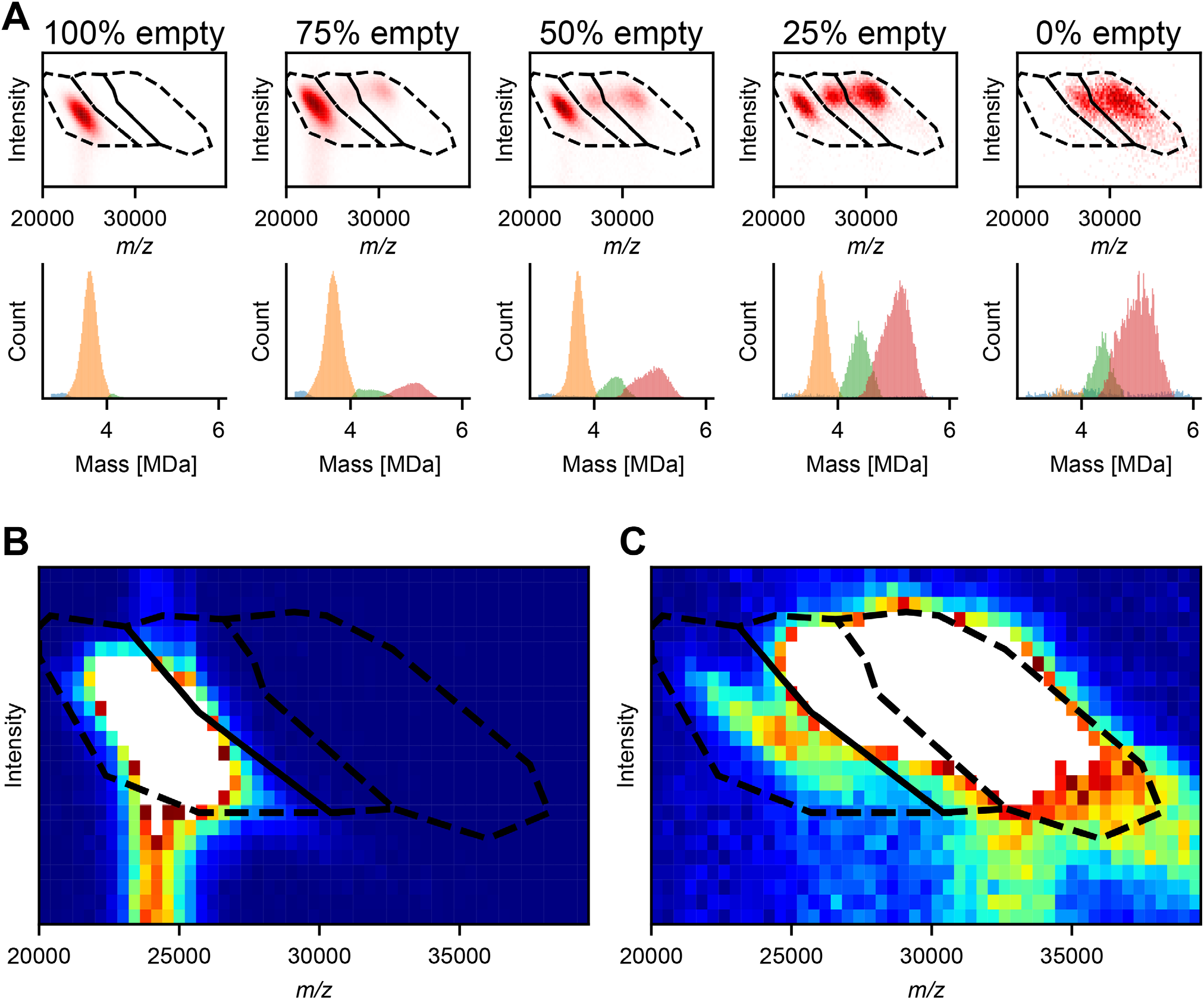
Quantification of distinct AAV6b particles at elevated pressure. **A)** Applied AAV quantification workflow on fractionated and mixed AAV particles at Xenon pressure setting 3. 2D histogram and corresponding mass histograms for a 20 minutes dataset of the seized particles mixed at defined ratios (amount of empty fraction indicated above). **B and C)** Pooled datasets of seized empty and full fraction displayed with the filtering masked used for quantification. The empty fraction (**B**) does not show any signs of filled particles, however we do observe a small amount of signal, leaking into the assigned margins for intermediate particles. For the filled particles (**C**) we can observe a clear, low-abundant distribution of empty particles, responsible for the systematic overestimation of the empty particles in Figure 4C, also highlighting the challenges associated with rAAV purification.

